# Multi-Omics Integration Predicts Cell-Specific Gene Regulatory Response and Rhizosphere Dynamics in Maize Root Fertilizer Treatment

**DOI:** 10.64898/2026.08.16.744839

**Authors:** Jade Horcoff, Anuradha Goswami, Bharat Mishra

## Abstract

Improving nitrogen use efficiency in maize (Zea mays) requires understanding how distinct root cell types and regulatory networks process fertilizer inputs. Given the current limited understanding of fertilizer-induced, cell-type-resolved maize roots and regulatory networks, computational biology frameworks are needed to model and predict how nutrient inputs are translated into transcriptional responses. Here, we integrated fertilizer-induced maize root bulk RNA-seq with reference atlases of single-cell RNA-seq and scATAC-seq to construct and predict a cell-specific regulome of the maize root under inorganic and mixed amendments. We demonstrate that inorganic fertilization induced stress associated and management pathways. Regulome analysis identified transcription factors (TF) from the AP2/ERF, NAC, HSF, and WRKY superfamilies that were preferentially active across root tissues. Deconvolution of the regulome onto single-cell atlases predicted core TF activity to the vascular cylinder and pith across both regimes, while mature cortex regulatory programs diverged. Construction of a gene regulatory network revealed that shared TF–target edges maintained the same regulatory orientation across fertilizer regimes. However, a small number of stress related TFs, including WRKY24, DREB1A, and NAC61, underwent a directional change between fertilization treatments. In silico knockout analysis predicted the activation targets for six of the seven regulators in their resident vascular/pith tissues, indicating the network behaves as a coherent, perturbable system. Additionally, soil metagenomic analysis showed that host soil microbial functions overlap with differentially expressed genes (DEGs) in shared functional categories, linking host regulome dynamics to rhizosphere processes. These findings and predictions suggest that the maize root regulome is spatially organized and dynamically reprogrammed by master regulators, predicting high-priority candidate nodes for engineering improved nutrient use efficiency.

## 1. Introduction

Maize (*Zea mays* L.) is among the most nitrogen-demanding cereal crops, and inorganic NPK fertilization remains the predominant strategy for sustaining the yields that support global food and feed supply. Yet nitrogen use efficiency (NUE) rarely exceeds 30–40% in maize cropping systems (Hirel, Le Gouis et al. 2007). Nitrogen that is not captured is lost through nitrate leaching into groundwater and surface water, nitrous oxide emissions, and ammonia volatilization. Fertilizer over-application increases costs for growers and is a major driver of agricultural eutrophication and greenhouse-gas emissions. Improving NUE is therefore a central objective of sustainable agriculture and molecular breeding programs, and requires a mechanistic understanding of how the root perceives and responds to fertilizer inputs (Hirel, Le Gouis et al. 2007).

The root is organized into concentric tissue layers, including the epidermis (with the lateral-root cap, LRC), cortex, endodermis and pericycle, which surround a central vascular cylinder (the stele) that contains xylem, phloem and pith. At the root apex, the quiescent center (QC) and surrounding meristematic initials sustain indeterminate growth (Hochholdinger and Tuberosa 2009). These cell types differ in their transport functions, hormonal environments, developmental trajectories, and nitrogen signaling is spatially partitioned among them. Nitrate uptake and long-distance signaling are concentrated in epidermal and vascular tissues, whereas nitrogen-dependent developmental decisions such as lateral-root initiation are executed in the pericycle (Zhang and Forde 1998, Gojon 2017). Accordingly, fertilizer responses and regulatory events differ substantially among these cell populations. Single-cell and single-nucleus genomics now make it possible to recover this spatial resolution. Recent single-cell RNA-sequencing (scRNA-seq) of maize root has produced annotated atlases spanning the full complement of root cell types and cell-type-resolved responses to nitrate (Ortiz-Ramírez, Guillotin et al. 2021, Li, Zhang et al. 2022). Additionally, single-cell ATAC-sequencing (scATAC-seq) has mapped the accessible *cis*-regulatory landscape of those same cell types, revealing where regulatory DNA is open to transcription factor binding (Marand, Chen et al. 2021). With current limitations in treatment-matched single-cell profiling, these reference atlases can be leveraged computationally for pseudobulk-based deconvolution of bulk RNA-Seq samples into cell-type proportions, and cell-type marker sets for most likely active DEGs.

Transcription factors (TFs) act as the control nodes that convert nutrient and stress signals into coordinated gene-expression programs. Several TF families are established regulators of drought and nutrient responses in the reference dicot *Arabidopsis thaliana* and have conserved co-orthologues in maize. The AP2/ERF superfamily is the largest plant-specific TF family, which integrates ethylene (ERFs) and abiotic-stress signaling (DREBs) to downstream transcription (Nakano, Suzuki et al. 2006, Licausi, Ohme-Takagi et al. 2013). NAC TFs govern secondary cell-wall biogenesis and nutrient-remobilization senescence (Uauy, Distelfeld et al. 2006, Mitsuda, Iwase et al. 2007). WRKY factors mediate nitrogen-responsive and immune-priming circuits, with several acting in the phloem in systemic nitrogen signaling (Rushton, Somssich et al. 2010). MYB, homeobox/HD, HSF/Sigma and TIFY/JAZ families contribute additional layers of control over metabolic, developmental, thermotolerance and Jasmonate (Chini, Fonseca et al. 2007, Dubos, Stracke et al. 2010). Gene-expression change alone cannot distinguish a directly acting regulator from a passively co-regulated gene; therefore, these TF families must be linked to their target genes and to the specific cell types to understand their regulatory impact during fertilization treatment.

The plant root functions within a rhizosphere microbiome whose composition and metabolic activity respond strongly to fertilization; the microbiome, in turn, influences plant nitrogen nutrition and hormone status. This plant–microbiome system is increasingly viewed as a holobiont whose combined genome shapes host phenotype (Vandenkoornhuyse, Quaiser et al. 2015). Convergent functional remodeling of the host transcriptome and associated microbiome in response to the same nutrient perturbation would indicate a co-regulated response system. However, such host–microbiome functional connections are rarely examined at the pathway level and are almost never resolved to specific host cell types within a single study.

To address these knowledge gaps, we integrated bulk RNA-seq profiling of maize roots under fertilizer regimes with reference single-cell transcriptomic (scRNA-seq) and chromatin accessibility (scATAC-seq) atlases, along with soil shotgun metagenomics data. By deconvolving cell-type-specific responses and building chromatin-supported gene regulatory networks (GRNs), we mapped the spatial partitioning of root nutrient responses across tissue layers. This framework revealed that AP2/ERF factors stay activated, whereas WRKY24, DREB1A, and NAC61 change the regulatory direction between inorganic and organic amendments. We validated the core regulatory nodes against independent nutrient-deficiency and single-cell nitrate datasets. Furthermore, we identified functional category alignment between host differentially expressed genes and rhizosphere microbial abundance. Taken together, this study provides a cell-type-resolved view of the maize root regulome under contrasting fertilization regimes and nominates concrete master regulator targets for engineering improved nutrient use efficiency in sustainable cereal crop production.

## 2. Materials and Methods

### 2.1 Public Dataset Retrieval and Treatment Conditions

Publicly available transcriptomic (GSA: CRA014161) China National Center for Bioinformation/Beijing Institute of Genomics, and metagenomic datasets were retrieved from the National Center for Biotechnology Information (NCBI) Sequence Read Archive (SRA) under the BioProject accession number PRJNA607213 for root responses to distinct fertilization regimes. ScRNA-seq dataset (GEO: GSE173087) (Ortiz-Ramírez, Guillotin et al. 2021) and chromatin accessibility profiling, the scATAC-seq dataset (GEO: GSE155178) (Marand, Chen et al. 2021), were downloaded from NCBI. Additional validation datasets were retrieved from NCBI GEO, including a nitrogen deficiency dataset (GSE125417) and an acute single-cell nitrate response dataset (GSE183171).

### 2.2 Bulk RNA-seq Data Analysis

Raw paired-end RNA-seq reads for six samples across three treatment groups (Control = CK, Inorganic = NPK, and Mixed = OMNPK) were processed using the *nf-core/rnaseq* pipeline (v3.18.0) executed via Nextflow with Singularity containers for reproducibility (Ewels, Peltzer et al. 2020). Read-quality trimming and adapter removal were performed with *fastp*, followed by ribosomal RNA depletion using *SortMeRNA*. Trimmed reads were aligned to the *Zea mays* B73 reference genome (GCF_902167145.1) using *STAR* in two-pass mode, with gene annotations obtained from the nuclear-chromosome-only GTF derived from the same assembly. Duplicate reads were marked using *Picard MarkDuplicates*, and alignment indices were generated in CSI format to accommodate the large chromosome sizes of the maize genome. Transcript- and gene-level quantification was performed with *Salmon* in alignment-based mode (via *STAR-Salmon*) and independently in quasi-mapping mode as a pseudo-aligner. Differential expression analysis was performed using *DESeq2* (v1.34.0). Genes with an adjusted *p-*value (<0.05) and an absolute log_2_fold change > 1 were identified as differentially expressed genes (DEGs).

### 2.3 Single-Cell Reference Atlas Integration and Cell-Type Deconvolution

Two published single-cell maize root atlases were used as orthogonal references. For transcriptional cell-type resolution, the scRNA-seq dataset was processed as a Seurat (v4) object retaining the original cell-type annotations (Hao, Hao et al. 2021). For chromatin accessibility profiling, the scATAC-seq dataset was loaded as a sparse binary accessible chromatin region (ACR) × nucleus matrix. Nucleus identities and cell-type assignments were obtained from the published supplemental barcode metrics table (Stuart, Srivastava et al. 2022). ACRs were represented as GenomicRanges coordinates anchored to the B73 reference genome AGPv4 (Lawrence, Huber et al. 2013). To associate ACRs with specific transcription factor (TF) loci, a gene-coordinate window was defined from 2,000 bp upstream to 500 bp downstream of each annotated transcription start site in AGPv4. TF LOC identifiers were resolved to AGPv4 locus coordinates, and overlapping ACRs were identified using GenomicRanges (Lawrence, Huber et al. 2013).

### 2.4 TF Activity Thresholding and Cross-Dataset Cell-Type Harmonization

TF activity in each cell type was quantified independently in each atlas. For scATAC-seq, a TF–cell-type pair was considered accessible if ≥20% of nuclei within that cell type contained at least one open ACR overlapping the TF gene window (pct_accessible ≥ 20). For scRNA-seq, a TF–cell-type pair was considered expressed if ≥15% of cells in that population expressed the gene at detectable levels (pct.exp ≥ 15), as calculated using Seurat’s DotPlot function applied to the 17 query TFs. Because the two atlases employ distinct cell-type nomenclatures, all cell types were mapped to a common set of nine harmonized biological groups: QC/Initials, Epidermis, Pericycle, Phloem, Xylem, Stele/Vasculature, Endodermis, Cortex, and Pith. Concordance of TF activity across the two modalities was assessed by identifying TFs whose active biological groups overlapped between datasets. The number of shared groups for each TF was used to rank TFs according to cross-dataset support.

### 2.5 Pseudobulk Signature Matrix Construction and Single- Cell Deconvolution

To infer the cell-type composition of bulk root RNA-seq samples, a pseudobulk reference signature matrix was constructed from the scRNA-seq atlas. Log-normalized expression values were retrieved from the data layer of the Seurat RNA assay, and mean expression was calculated across all cells of each annotated cell type, excluding the "Uncharacterized" population. Bulk gene symbols from the Salmon-quantified count matrix were bridged to AGPv4 Zm00001d gene identifiers using NCBI gene info for maize, retaining a single Zm00001d locus per symbol where duplicate mappings occurred. Bulk counts were first normalized to counts-per-million (CPM) and subsequently log1p-transformed to match the scale of the scRNA-seq reference. Deconvolution was restricted to genes present in both the bulk and signature matrices. To improve the signal-to-noise ratio, the 5,000 genes with the greatest cross-cell-type variance in the signature matrix were selected as informative features. Cell-type proportion estimation was performed independently for each bulk sample using non-negative least squares (NNLS) regression, as implemented in the R *nnls* package, with the signature matrix as the design matrix. Raw NNLS coefficients were normalized to sum to one, yielding fractional cell-type proportions for each sample. Deep neural-network (DNN) ensemble was also implemented with Scaden architecture trained on pseudobulk mixtures simulated from the same atlas (Menden, Marouf et al. 2020). Estimated cell-type proportions were compared across fertilizer treatments.

### 2.6 DEG-Based Cell-Type Enrichment Analysis

As a complementary approach, cell-type enrichment of treatment-responsive genes was assessed by hypergeometric testing. Cell-type marker specificity was quantified for each gene as the ratio of its mean expression in a given cell type to its mean expression across all cell types (with a pseudocount of 0.01). For each cell type, the top 200 genes by specificity score were designated as the representative marker set. For each treatment contrast, DESeq2-significant differentially expressed genes (adjusted p < 0.05) with available Zm00001d identifiers were used as the query set. Hypergeometric enrichment of each cell-type marker set within the DEG list was computed with the full set of genes detected in the scRNA-seq atlas as the background universe. This analysis identified the root cell populations that were most transcriptionally responsive to each fertilizer treatment.

### 2.7 Chromatin-Supported Gene Regulatory Network (GRN) Construction

To construct a high-confidence, multi-tiered gene regulatory network (GRN), transcription factor (TF) binding predictions were combined with single-cell chromatin accessibility and bulk expression dynamics. Transcription factor binding sites (TFBS) were mapped across promoter regions (2KB) relative to transcription start sites of maize genes using position weight matrices (PWMs) from the JASPAR (Castro-Mondragon, Riudavets-Puig et al. 2022) and PlantTFDB databases (Jin, Tian et al. 2017). Predicted binding sites were filtered against open chromatin regions defined by root scATAC-seq peak calls to ensure biological relevance. Co-expression relationships between accessible TFs and downstream DEGs were computed using GENIE3 (Chen and Mar 2018) and ARACNE (Margolin, Nemenman et al. 2006). TF–target edges were classified as activating or repressing based on Directional Pearson Correlation (r). Edges exhibiting consistent regulatory signs across both NPK and OMNPK regimes were designated as the sign-invariant network core.

### 2.8 Soil Metagenomics and Functional Analysis

Metagenomic sequence processing, assembly, binning, and taxonomic profiling were executed reproducibly using the nf-core/mag pipeline managed via Nextflow and Singularity (Krakau, Straub et al. 2022). Initial preprocessing included contaminant filtering against PhiX and Lambda phage references, digital read-depth normalization using BBNorm, and ancient DNA damage profiling to account for degraded fragments. High-quality reads were assembled *de novo* using MetaSPAdes (--meta) and subsequently mapped back to contigs using Bowtie2 in --very-sensitive mode (binning_map_mode = own). Contig bins were integrated and refined using DAS Tool (score threshold: 0.30), with bin completeness and quality assessed using BUSCO (bacteria_odb10) alongside raw CheckM diagnostics. Finally, taxonomic classification of reads and assembly contigs was performed using a multi-database consensus approach leveraging Kraken2, Centrifuge, and CAT (Contig Annotation Tool). Protein-coding genes were predicted using Prodigal (v2.6.3) and clustered at 95% sequence identity using CD-HIT. Functional annotation was conducted by aligning predicted amino acid sequences against Gene Ontology (GO), KEGG Orthology (KO), and eggNOG databases using DIAMOND (v2.0.13). Differential abundance of functional categories between fertilizer regimes was evaluated using DESeq2 on mapped functional feature counts.

### 2.9 Statistical Analyses, Enrichment Controls, and TF Family Size Normalization

All statistical computations were performed in R (v4.2.1). Gene Ontology and pathway enrichment analyses were conducted using hypergeometric tests with Benjamini-Hochberg false discovery rate (FDR) correction (FDR < 0.05). To avoid artificial overrepresentation bias when identifying key regulator superfamilies (e.g., AP2/ERF, WRKY, NAC), TF family enrichment was normalized against total family representation within the B73 reference genome:

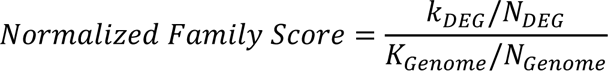

where *k_DEG_* is the number of DEGs in a given TF family, *N_DEG_* is the total number of TF DEGs, *K_Genome_* is the total number of members in that TF family across the genome, and *N_Genome_* is the total number of TFs in the genome. TF families were classified as enriched only when the normalized score exceeded baseline genome expectations.

### 2.10 In Silico Transcription-Factor Knockout Analysis

To extend the correlative regulome into a predictive, perturbable model, the seven-regulator base network was propagated through the single-cell atlas using CellOracle v0.20.0 (Kamimoto, Stringa et al. 2023). The single-cell atlas was imported as an AnnData object; the base gene regulatory network was supplied directly as the study’s own TF–target edge set (7 TFs, 984 edges) rather than from a motif scan. CellOracle’s raw-count layer was reconstituted as the linearized normalized expression rounded to integers. Following PCA (37 components) and balanced k-nearest-neighbor imputation (k = 365), cell-type-specific GRN models were fitted using bagged ridge regression for all 22 cell types, and coefficient matrices for signal propagation were estimated (α = 10). In silico knockout of each hub regulator was simulated by setting its expression to zero and propagating the resulting effect through the fitted network for three iterations. For each cell type, the perturbation effect was quantified as the mean signed and absolute shift (simulated − imputed expression) across that regulator’s activation and repression targets.

### 2.11 Diffusion Pseudotime and Trajectory-Resolved Expression

To place the regulome in a developmental context, diffusion pseudotime (DPT) was computed for the single-cell reference atlas (Haghverdi, Buttner et al. 2016). Log-normalized expression of the 14,591 annotated cells was reduced to the 2,000 most variable genes, scaled, and embedded by principal component analysis (30 components); a k-nearest-neighbor graph (k = 30) and diffusion map (15 components) were built with Scanpy (Wolf, Angerer et al. 2018). Pseudotime was rooted at the quiescent center (the atlas cell nearest the QC centroid in diffusion space) and computed with Scanpy’s dpt function. The ordering was validated against the canonical root developmental sequence (quiescent center -> initials -> early/meristematic -> mature/differentiated tissues) using Spearman’s correlation between mean per-cell-type pseudotime and developmental-stage rank. Predicted regulon activity was then plotted along the pseudotime axis for each hub regulator and treatment. For the trajectory-resolved transcriptome, the union of inorganic and mixed treatment DEGs present in the atlas (1,848 genes) was averaged across 30 equal-cell pseudotime bins, z-scored by gene. The resulting expression profiles were hierarchically clustered (Ward linkage) into co-expression modules ordered by peak pseudotime. Each module was then tested for Gene Ontology enrichment against the genome-wide maize gene2go background (hypergeometric test, Benjamini–Hochberg correction).

### 2.12 Software and Reproducibility

Analyses were run in R (≥ 4.3; Seurat and SeuratObject for single-cell handling, Matrix, data. table, GenomicRanges, dplyr, ggplot2, patchwork, and nnls for the linear deconvolution) and Python (≥ 3.10). Bulk differential expression used the deposited DESeq2 tables and was independently reproduced with DESeq2 for the two-treatment volcano analysis. Single-cell processing, diffusion pseudotime analysis, and imputation were performed using Scanpy and AnnData. In silico transcription-factor knockouts were performed using CellOracle. Deep-ensemble deconvolution was reimplemented from the Scaden architecture with scikit-learn (MLPRegressor). Soil-metagenome taxonomic classification was performed using Kraken2 and aggregated from the phylum to genus level. Hypergeometric tests and Benjamini–Hochberg corrections were performed using SciPy and statsmodels. Gene Ontology background sets were derived from the NCBI Zea mays gene2go annotation, and gene identifiers were bridged across genome builds throughout using the NCBI gene_info LocusTag↔Symbol table (Zm00001d↔LOC/Entrez). Figures were generated with matplotlib.

## 3. Results

### 3.1 Fertilizer treatments induce large, directionally biased transcriptional responses

Both the inorganic (NPK) and mixed organic/inorganic (OMNPK) amendments drove large, directionally biased transcriptional responses relative to control roots. Each regime was sampled with two biological replicates; therefore, we assessed the robustness of the DEG results given the limited replication before using them in subsequent analyses. Principal-component analysis of the six libraries separated fertilized from control roots along PC1 (57.4% of variance), with replicates of each regime clustering together and a between-group/within-group distance ratio of 1.69 (Figure S1). An independent effect-size estimate computed directly from the Salmon count matrix reproduced the deposited DESeq2 log₂ fold changes with high fidelity (Pearson *r* = 0.96 across 27,763 genes). Inorganic NPK fertilization displayed a large, repression-biased transcriptional response. A total of 1,235 genes were differentially expressed relative to control roots (FDR < 0.05; 492 upregulated, 743 downregulated; Figure 1A). The mixed (OMNPK) contrast yielded a comparable set of 1,278 DEGs (555 upregulated, 723 downregulated; Figure 1B) using the same DESeq2 significance criteria (Table S1).

**Figure 1.**
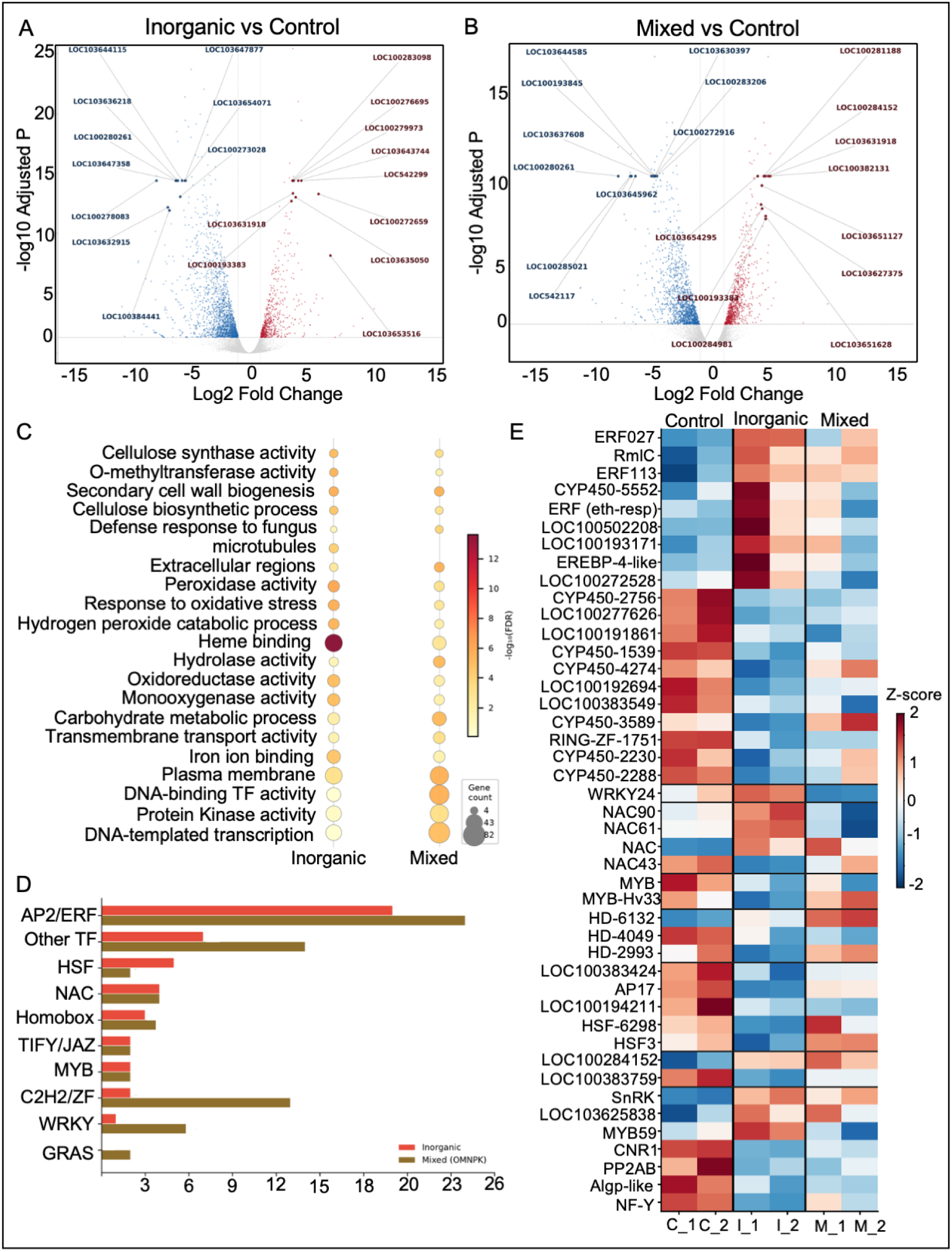
Differential expression under the two fertilizer regimes. (**A**) Inorganic (NPK) vs control; (**B**) mixed (OMNPK) vs control. Each point represents a gene; red = up-regulated, blue = down-regulated (adjusted *p* < 0.05 and |log₂FC| ≥ 1), grey = not significant. Dashed line, *p* = 0.05; dotted lines, |log₂FC| = 1. (**C**) Gene Ontology over-representation of the inorganic and mixed DEG sets, computed identically against the NCBI Zea mays gene2go annotation (FDR < 0.05) in at least one regime and among the top by FDR are shown; dot size encodes fold-enrichment and color encodes *−log₁₀* FDR. Both regimes share an oxidative/redox and secondary-cell-wall core; the mixed regime additionally enriches transcriptional-regulation, carbohydrate-metabolism and defense-response terms. (**D**) Number of maize DEGs per TF family under the inorganic and mixed contrasts. AP2/ERF contributes the most DEGs under both regimes; WRKY and C2H2/zinc-finger families expand markedly under the mixed treatment. (**E**). Expression of the 44 differentially expressed transcription factors across the bulk RNA-seq libraries, z-scored per gene on log₂ counts and grouped by TF family.

To compare functional enrichment between the two fertilizer regimes, we re-ran GO over-representation analysis for both DEG sets against a single genome-wide annotation (Figure 1C, hypergeometric test, BH-corrected *P* <0.05). Both regimes converged on a shared functional core of oxidative/redox metabolism (peroxidase activity, response to oxidative stress, hydrogen peroxide catabolism), monooxygenase and heme-binding activity, and secondary cell-wall/cellulose biogenesis. Interestingly, heme binding, peroxidase, and H₂O₂-catabolic terms were more strongly and significantly enriched under inorganic NPK, whereas the mixed regime showed stronger enrichment of transcriptional-regulation terms (DNA-binding transcription factor activity, regulation of transcription), carbohydrate/O-glycosyl-hydrolase metabolism, and, distinctively, regulation of defense response to the GO-level counterpart of the Jasmonate/pathogenesis-related signature seen among the mixed top DEGs.

To explore the regulator’s activity, we identified 48 and 77 DEGs as transcription factors responsive to inorganic and mixed treatments, respectively. Interestingly, eight TF families, including AP2/ERF (ERF34, ERF027, ERF113, EREBP-4-like), NAC-family (NAC61, NAC90, ANAC053-like), HSF/Sigma (HSF3, HSF1), Homeobox/HD (ATHB-40, HD1, HOX22), and WRKY (WRKY24), are most enriched in the fertilizer-treated maize roots (Figure 1D, Table S1). AP2/ERF contributed the largest share, 20 of the 44 DEGs (9 up, 11 down), including ERF027, ERF113, and EREBP-4-like (Figure S1D), while the HSF/Sigma factors are uniformly high in control and repressed under both fertilizers. The dual up/down pattern indicates a bifurcated response: suppression of oxidative-stress cytochrome P450 isoforms alongside induction of ethylene-mediated growth and nutrient-signaling programs. NAC-family DEGs (3 upregulated, 1 downregulated) were consistent with the secondary cell-wall enrichment. All five HSF/Sigma DEGs were down-regulated, consistent with reduced thermal stress load or feedback on plastid gene expression. The single WRKY member (WRKY24) links to nitrogen-responsive and immune-priming circuits. Overall, these gene-level patterns corroborated the GO-enrichment signal of redox rebalancing and secondary-cell-wall/secondary-metabolism reconfiguration under inorganic NPK fertilization.

### 3.2 Predicted Cellular localization of fertilization response

Deconvolving bulk RNA-Seq libraries against the 21-cell-type signature with a single validated NNLS pipeline (see Methods) placed the response in specific root tissue layers. Both fertilizer regimes drove a coordinated shift toward central-vascular and stele programs and away from epidermal differentiation, but each carried a distinct cortical signature (Figure 2A, Figure S2, Table S2). Under inorganic NPK, the strongest gains occurred in QC (+0.044), Stele (+0.037), and Early Stele (+0.016), while Mature Outer Cortex I (−0.035), Xylem (−0.026), and Epidermis/LRC (−0.020) showed reductions. The mixed regime produced an even larger gain in Early Stele (+0.049) and a comparable gain in Stele (+0.035). However, the cortical response diverged sharply, with Mature Outer Cortex I increase (+0.065), while Mature Cortex II showed the largest single loss in either treatment (−0.074). Thus, both fertilizers converge on a stele-biased composition, whereas the mature-cortex compartment is remodeled in opposite directions, predicting increase in cell counts under the mixed regime and decrease in cell counts under inorganic NPK.

**Figure 2:**
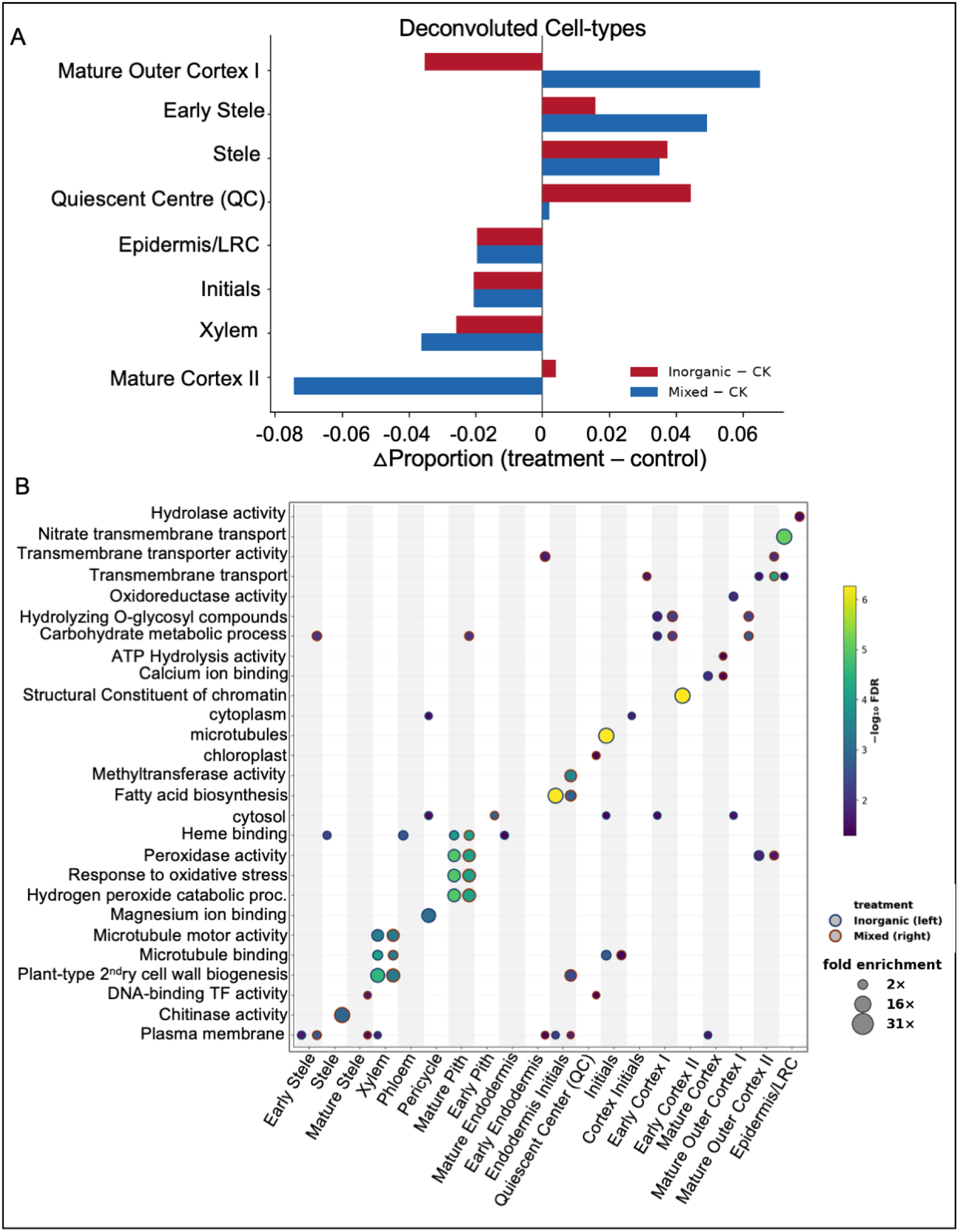
Deconvoluted Cellular localization of root fertilizer response. (**A**) Fertilizer-driven shifts in estimated cell-type proportions for the inorganic and mixed regimes relative to control (NNLS); bars show Δ proportion (treatment − control) for each pair. (**B**) Cell-type-resolved Gene Ontology enrichment of the fertilizer response. Each treatment’s DEGs were assigned to the cell type in which their deconvolution-signature weight is highest, and GO enrichment was computed per cell type against a genome-wide maize gene2go background (hypergeometric test, BH-corrected). Inorganic (left dot) and mixed (right dot) are shown side by side at each cell type; dot colour is −log₁₀ FDR and size is fold enrichment. Vascular and cortex programs dominate secondary cell wall biogenesis in Xylem, peroxidase/oxidative-stress terms in Mature Pith, nitrate transport in Epidermis/LRC with a small number of treatment-specific terms (e.g. chitinase activity in Stele under the mixed regime).

To test whether this linear deconvolution under-resolves low-abundance tissues, we re-estimated composition using a deep neural network (DNN) ensemble based on the Scaden architecture trained on pseudobulk mixtures simulated from the same atlas (Figure S2C, Table S2). The two methods agreed on the abundant, well-defined populations (Pearson *r* = 0.47–0.70) but differed sharply in resolution: NNLS assigned exactly zero proportion to 13 of the 21 cell types across all six libraries, concentrating roughly half of every library into Xylem and resolving a mean of 6.0 cell types per library, whereas the deep model assigned non-zero proportion to nearly every cell type. Reassuringly, both methods detected the low-abundance quiescent center (QC) and agreed that it expands under fertilization, and both recovered the stele-biased shift. A complementary hypergeometric enrichment analysis of DEGs in cell-type marker sets was performed for both contrasts (Figure S2D). The inorganic DEGs were significantly enriched in 12 cell types and the mixed-regime DEGs in 13, including Mature Pith (inorganic fold = 3.14, FDR = 9.8×10⁻⁷; mixed fold = 2.82, FDR = 2.5×10⁻⁵) and shared strong enrichment in Phloem, Stele, Xylem, and Mature Stele. This clearest contrast-specific signal also mirrors the deconvolution, including the mature outer/other cortex classes (Mature Outer Cortex I/II, Mature Cortex I/II) and more strongly enriched under the mixed regime. Resolving the functional response within each predicted cell type showed that the two regimes engage overlapping but distinct programs in the same tissues. Secondary cell wall biogenesis dominates in Xylem, peroxidase/oxidative-stress terms were observed in Mature Pith, and nitrate transport terms were observed in Epidermis/LRC. A small number of treatment-specific terms, such as chitinase activity in Stele under the mixed regime (Figure 2B, Table S2). Taken together, both fertilizers reprogram the same vascular-and-pith core, with the mixed regime additionally engaging the mature cortex.

### 3.3 A cell-type-resolved transcription-factor regulome

By integrating bulk RNA-Seq, scRNA-Seq pseudobulk co-expression, scATAC accessibility, and cross-modal concordance, we inferred a GRN for seven prioritized TFs and assigned them to three confidence tiers (1,2, and 3) (Figure S3, Table S3). High-confidence Tier 1 TFs include EREBP-4-like, with 289 candidate targets (|r| ≥ 0.50), and high-confidence chromatin-supported edges (|r| ≥ 0.80) in Epidermis/LRC, Pericycle, Phloem, and Xylem. Most of the EREBP-4-like targets are enriched in cytochrome P450 activity, iron-ion binding, and oxidoreductase activity. Another Tier 1 TF, the ethylene-responsive ERF, is scATAC-active in Pericycle, Phloem, and Xylem and scRNA-confirmed in six cell types, with three shared vascular cell types providing multimodal support; its targets are enriched for ethylene-activated signaling (FDR = 2.9×10⁻³). The medium-confidence Tier 2 TFs include WRKY24, which is active exclusively in phloem in both modalities, consistent with a phloem-restricted nitrogen-sensing/immune-priming role. Another Tier-2 TF, ERF113, shows scATAC accessibility across four cell types but non-overlapping scRNA expression, suggesting post-transcriptional regulation. NAC61 has robust scATAC support in four cell types but no cognate scRNA enrichment, possibly reflecting low-level or protein-level activity. The Tier 3 TFs are contextual and include NAC, which is broadly scRNA-active across seven cell types but lacks ATAC support. NAC’s 227 targets are enriched for chromatin structural components, suggesting protein–protein-mediated regulation. Another Tier 3 TF, DREB1A, the maize orthologue of cold/drought DREB1 factors, is scRNA-active in five cell types without ATAC support; its 88 targets are enriched for oxidative-stress response, H₂O₂ catabolism, and chitin catabolism.

To predict where each regulon is engaged, we scored each TF’s targets following the logic of expression-based activity methods (VIPER/AUCell). Each target’s treatment log₂FC was signed by its edge mode of regulation and distributed across the 21 cell types in proportion to that target’s atlas expression share. The resulting activity is spatially structured and reconfigures between regimes (Figure S3A-B). The ethylene-responsive ERF shifts from suppressed in all 21 cell types under inorganic NPK to active in 18 under the mixed regime. Conversely, the vascular-biased activity of EREBP-4-like and DREB1A, which dominates the inorganic response, collapses under the mixed amendment, while WRKY24, NAC, NAC61 and ERF113 gain broad activity that extends into the ground tissue (Figure S3C-D and S4). WRKY24 is transcriptionally down-regulated under the mixed regime yet shows broader predicted target-program activity, consistent with post-transcriptional or protein-level control.

To test whether that wiring behaves as a causal, perturbable system, we subjected the inferred regulome to in silico perturbation. Using CellOracle, we propagated the seven-regulator base network across the single-cell atlas and made each regulator perturbable by setting its expression to zero. We then propagated the effect through the fitted cell-type-specific networks to predict how its targets respond to its loss. The knockouts behaved as a coherent regulatory system rather than as a set of independent correlations (Figure 3A). For six of the seven regulators, in silico knockout collapsed the large majority of that factor’s activation targets EREBP-4-like 97.5%, the ethylene-responsive ERF 90.8%, WRKY24 100%, ERF113 92.7%, NAC 99.1%, and DREB1A 82.6% of activation targets down-regulated on knockout while de-repressing most of its repression targets (82.8–100%) (Table S3). The perturbation was tissue-structured and matched each regulator’s inferred niche. The EREBP-4-like knockout had its strongest effect in the Early Stele, the WRKY24 knockout in the Stele, and the NAC knockout in the Mature Pith (Figure 3B, Table S3). When we restricted EREBP-4-like’s targets to its activation edges, the knockout reduced their expression in vascular cells by a mean of −0.19 (96.3% of activation targets down), consistent with EREBP-4-like acting as a vascular activator of its cytochrome P450/oxidoreductase target module. Visualization of these knockouts as perturbation vector fields on the atlas embedding (Figure 3C-I) showed that each regulator drives a distinct, spatially coherent predicted cell-state shift rather than uniform noise.

**Figure 3:**
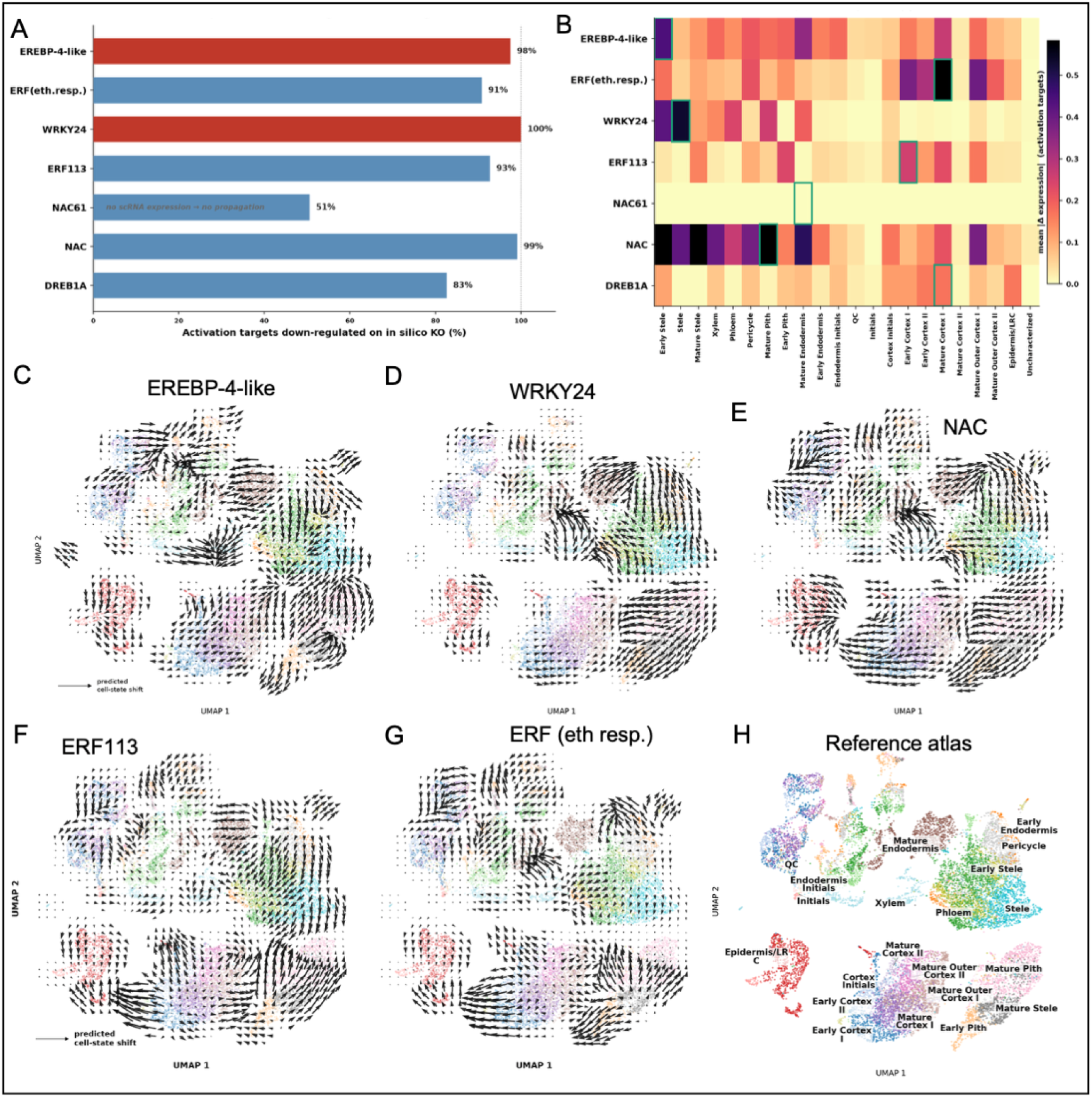
Regulome *in-Silico* Perturbation. **(A)** Percentage of each regulator’s activation targets that are down-regulated when the regulator is knocked out, with the two named hubs (*EREBP-4-like*, *WRKY24*) highlighted; the dotted line marks 100%. *NAC61* (51%) has chromatin accessibility but negligible single-cell expression, so its regulon cannot propagate. (**B**) Predicted perturbation magnitude (mean |Δ expression| over activation targets) for each regulator across the 22 cell types, ordered vascular to cortex. The strongest-effect cell type is boxed in green for each regulator. (**C-G**). Predicted direction of cell-state shift on in silico knockout, shown as CellOracle perturbation vector fields (quiver) on the single-cell UMAP embedding for hub regulators (EREBP-4-like, WRKY24, NAC, ERF113, and ERF (eth resp), and the single cell reference atlas (**I**). Each arrow shows the direction and magnitude in which CellOracle predicts a cell’s transcriptional state would shift if the regulator were removed. A coherent, long-arrow field localized over a tissue means the regulator actively maintains that tissue’s expression program there, while a flat field (e.g. NAC61) means the model predicts no dependence.

### 3.4 Integrating the regulatory model with the measured response

Placing each regulator’s CellOracle coefficients and predicted knockout magnitudes alongside the measured DESeq2 fold changes of the same targets under each fertilizer regime resolves the model and the data into one per-regulator view (Figure 4, Figure S5). We identified three consistent features across regulators. First, the CellOracle regulatory coefficient recovered the co-expression-inferred activation/repression sign for 323 of 359 shared-core edges (90%), indicating that the perturbable model and the correlation network agree on edge polarity. Second, within each shared core, the inorganic and mixed fold-change columns largely agree in sign, i.e 342 of the 359 shared targets (95%) move in the same direction under both regimes, complementing the edge-direction sign conservation, while the treatment-unique targets show a fold change only in their own regime (Table S3). Third, the regulator’s activity exposes the directional regulation, EREBP-4-like and NAC stay induced under both regimes, whereas WRKY24 changes from induced under inorganic NPK to strongly repressed under the mixed amendment, along with NAC61 and DREB1A. These results demonstrate that the predicted regulatory logic (edge sign and target membership) is largely conserved across regimes.

**Figure 4:**
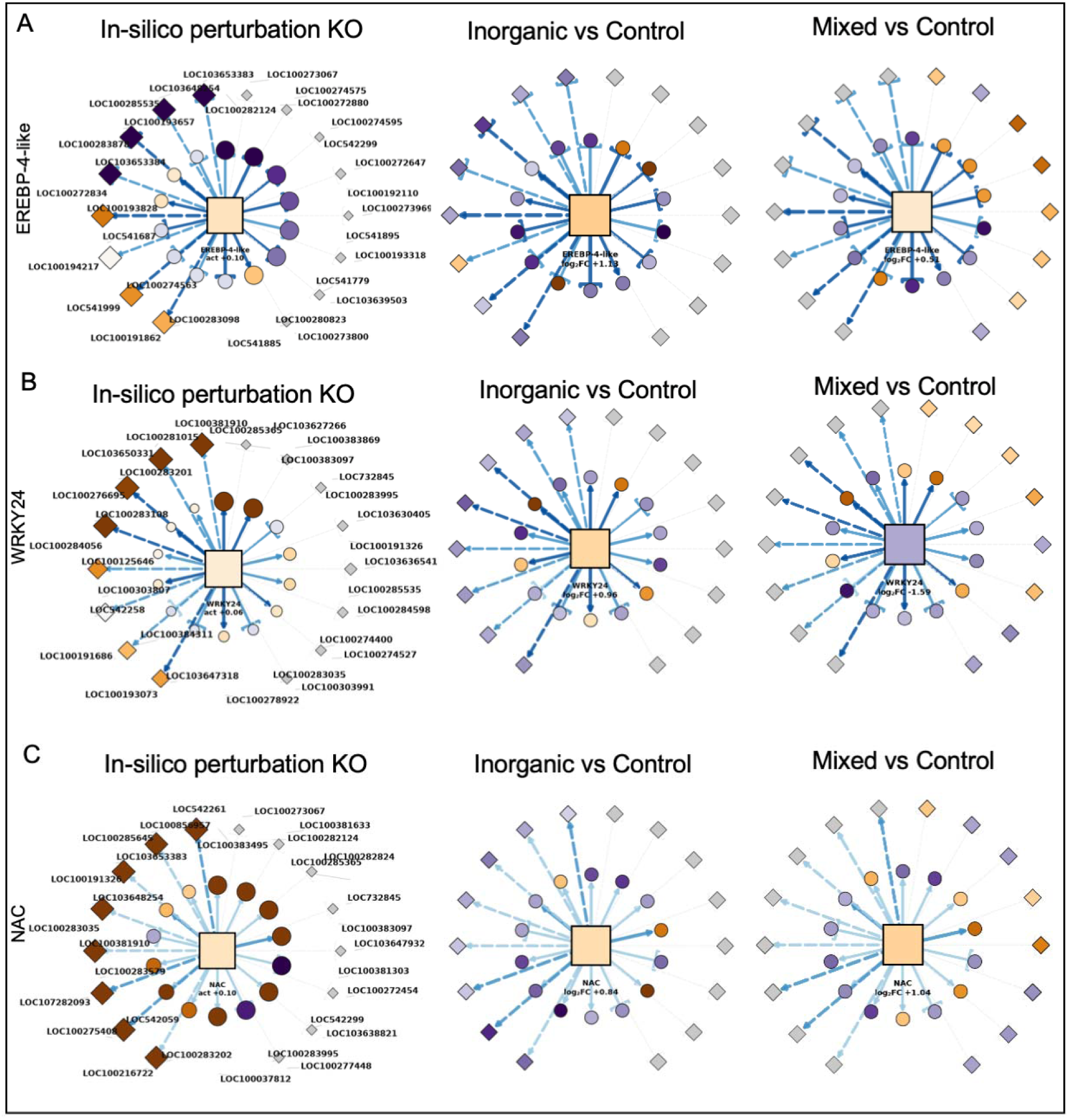
Fertilizer impacted GRN regulome. Per-regulator regulome for the three hub regulators (EREBP-4-like (A), WRKY24 (B), and NAC (C)). Each row is one regulator; the three columns show the CellOracle regulatory model (node color = regulatory coefficient, node size = in silico knockout Δ), the inorganic response and the mixed response (node color = target DESeq2 log₂ fold-change). Circles are shared-core targets, diamonds are treatment-unique targets; solid edges denote shared and dashed edges treatment-unique target links; arrowheads mark activation and bars repression; edge color encodes network confidence and edge width the co-expression |r|. The central square is the regulator, colored by its own activity in that column (mean predicted regulon activity for CellOracle, DESeq2 log₂FC for the two treatments). Node positions are matched across the three columns and gene identifiers are labelled once, on the CellOracle panel. Orange = activation/up-regulated, purple = repression/down-regulated, grey = unscored or absent in that view.

Finally, to determine whether the fertilizer-responsive regulome is organized along root development, a diffusion pseudotime was derived directly from the reference atlas and rooted at the quiescent center (Figure 5A). The ordering recovered the expected developmental axis: mean per-cell-type pseudotime correlated with the canonical stage sequence (Spearman ρ = 0.58, *P* = 0.006), with the quiescent center and initials at the origin and mature vascular and epidermal tissues at the late end (Figure 5B). Hierarchical clustering of the 1,848 union DEGs across pseudotime bins resolved six co-expression modules with distinct temporal peaks whose Gene Ontology signatures track developmental progression. Early modules were enriched for cell-division and cell-wall machinery (microtubule, vesicle coat, secondary cell wall biogenesis), whereas late modules were enriched for nitrate transport, chromatin components, and cellulose synthase (Figure S6). Predicted activity of the seven hub regulators varied along this axis rather than being uniform, and the two fertilizer regimes diverged across the pseudotime (Figure 5C-D). Because the atlas is a snapshot of many co-existing mature tissues rather than a single continuous lineage, the pseudotime spread is compressed, and the trajectory signal and these analyses are presented as predictive developmental context for the regulome.

**Figure 5:**
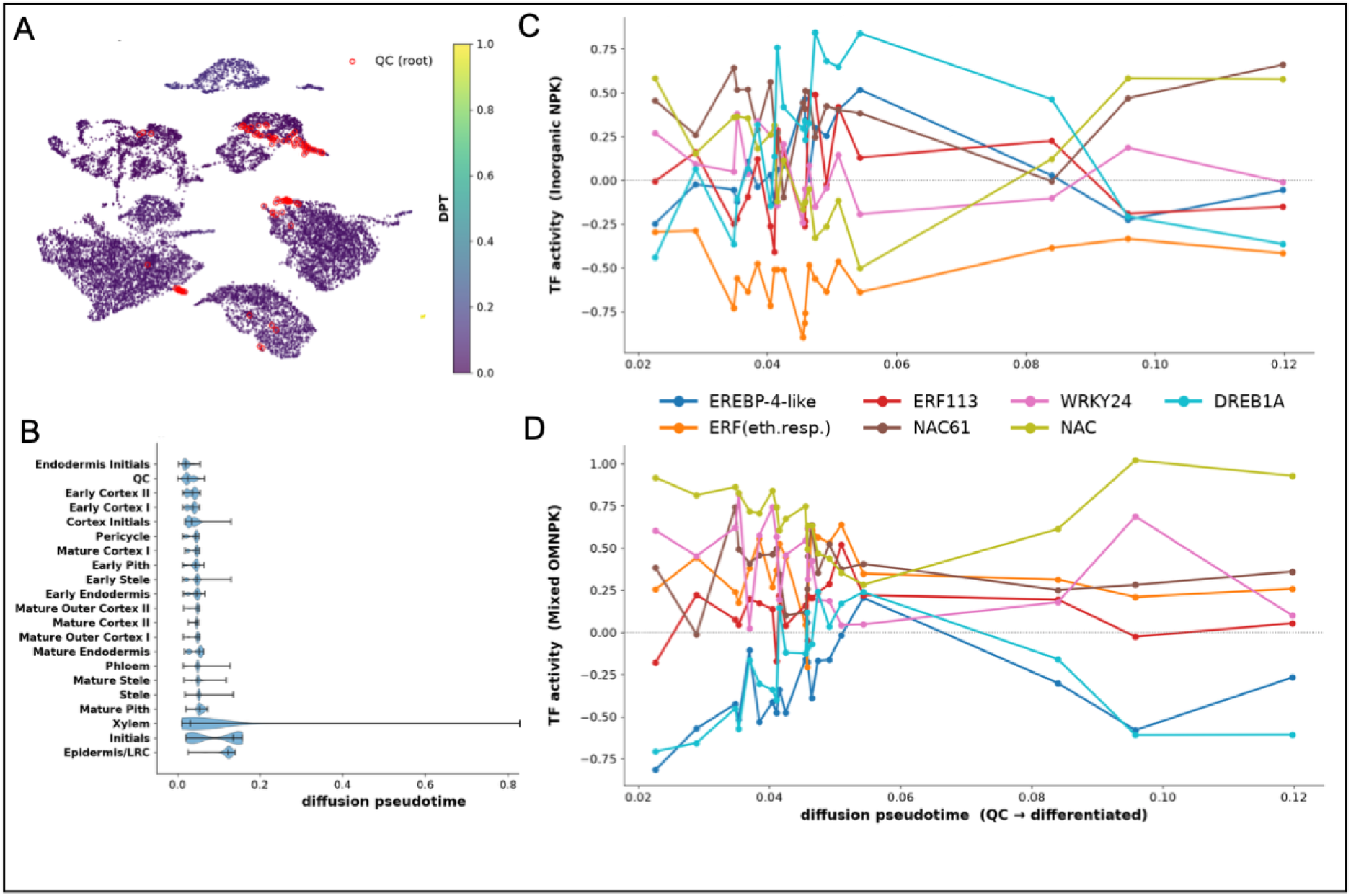
Diffusion pseudotime of the single-cell atlas and regulome activity. (**A**) Atlas UMAP (14,591 nuclei) colored by QC-rooted diffusion pseudotime, with quiescent-Centre cells outlined. (**B**) Pseudotime distribution per cell type, confirming the QC/initials root and the differentiated tissues (Epidermis/LRC, mature vascular) at the late end (Spearman ρ = 0.58 between mean per-cell-type pseudotime and canonical developmental stage, p = 0.006). (**C, D**) Predicted activity of the seven hub regulators along the diffusion pseudotime under inorganic and mixed fertilizer.

### 3.5 Fertilizer-Impacted Rhizosphere microbiome

Soil shotgun metagenomics profiled the rhizosphere community accompanying the fertilizer treatments. Taxonomic classification identified 812 genera at the genus level, and the two fertilized regimes were sharply less even than the control. The unfertilized community was diverse and *Streptomyces*-, *Pseudomonas*-, and *Burkholderia*-rich (Shannon H′ = 4.97; top genus 9.4%), whereas both fertilized communities were dominated by *Bacillus.* Its relative abundance increased from 0.5% of classified reads in the control to 70% under the organic amendment and 89% under inorganic NPK, accompanied by a corresponding collapse in genus-level diversity (H′ = 2.13 organic, 0.84 inorganic (Figure 6A-B, Table S4). This *Bacillus* enrichment under fertilization is consistent across the two high-depth libraries (>4 million classified reads for control and inorganic) rather than being a low-coverage artefact. Functional pathway annotation of the assembled metagenomes yielded broad repertoires of 2,265 (inorganic) and 1,986 (organic) pathway categories, dominated by transport (ABC/permease), transcriptional regulation, and amino acid/peptide metabolism (Figure 6C-D, Table S4). In a like-for-like comparison, the organic amendment showed uniformly higher, though modest, enrichment scores across nearly every functional category. The pathways most divergent between regimes were consistent with a microbial iron-scavenging emphasis under organic inputs, including higher iron-uptake permease, the enterobactin exporter *EntS,* and dipeptide transport. IN contrast, recombinase and two-component sensor functions were higher under inorganic fertilization (Figure 6D). The metagenome consisted of a single assembled sample per treatment and is therefore used here only to characterize the taxonomic and functional context of the host response, not to make replicated microbial comparisons.

**Figure 6:**
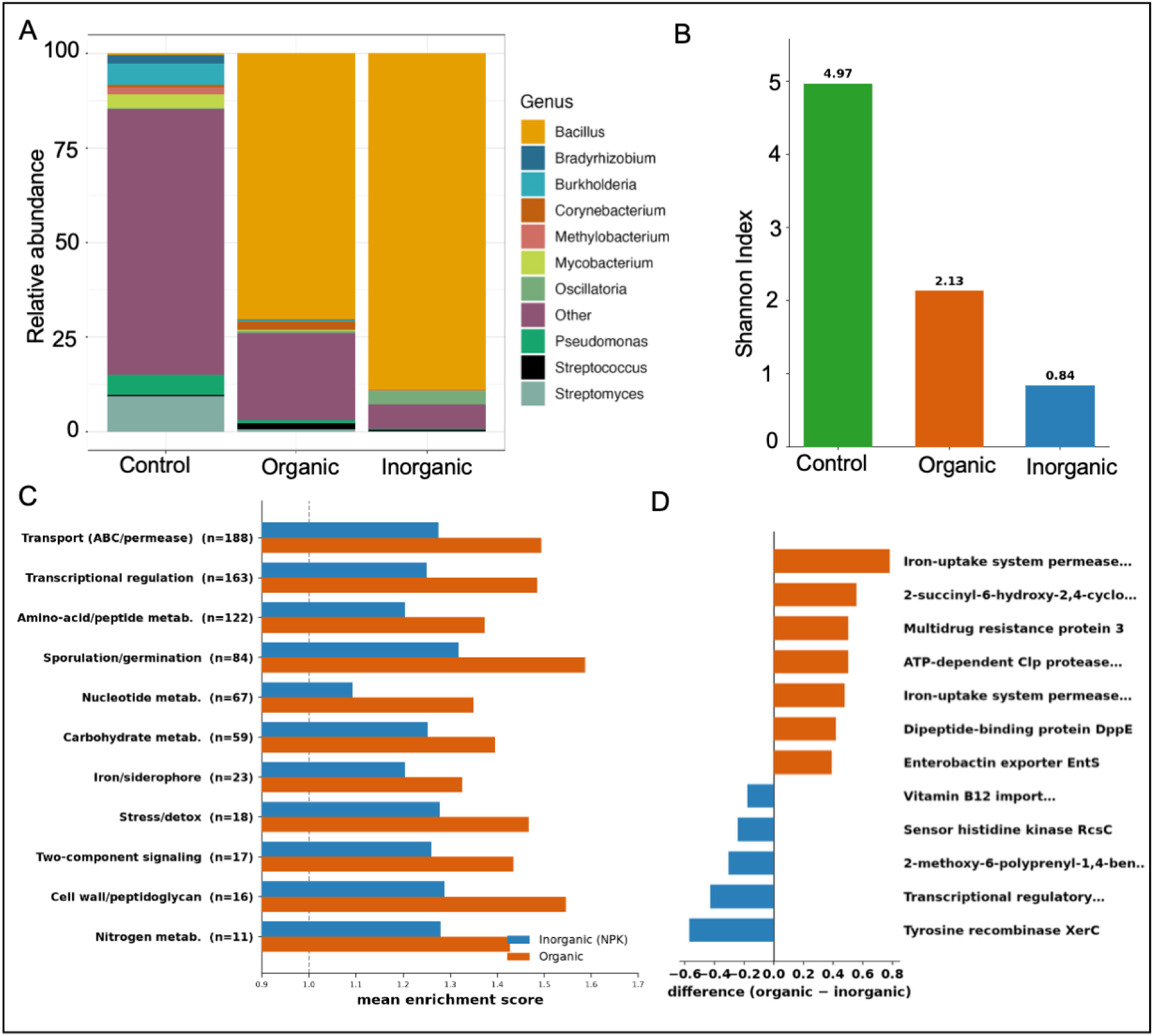
Rhizosphere composition and diversity by fertilizer regime. (**A**) Relative abundance of the ten most abundant genera (remaining 802 genera pooled as "Other") in the control, organic and inorganic metagenomes. (**B**) Shannon diversity computed on the genus-level relative abundances, showing a progressive collapse from the diverse control community to the Bacillus-dominated fertilized communities. (**C**) Mean functional-enrichment score per pathway category for each treatment (dashed line = no enrichment, score 1.0). (**D**) The individual pathways most divergent between regimes (enrichment-score difference, organic − inorganic; pathways with ≥ 5 assigned genes), showing an iron-scavenging/transport emphasis under organic inputs.

## 4. Discussion

By resolving bulk transcriptomic averages into a cell-type-structured, chromatin-grounded network, this study clarifies the precise molecular and spatial mechanisms by which maize roots sense and adapt to fertilizer inputs. Integrating bulk transcriptomics, single-cell RNA-seq (scRNA-seq), single-cell ATAC-seq (scATAC-seq), and *in silico* perturbation modeling reveals that the maize root transcriptional response to fertilization is directed by a spatially structured, AP2/ERF-dominated regulome. Rather than acting as a uniform global shift, nutrient amendment triggers targeted transcriptional reprogramming centered within specific tissue layers and controlled by a discrete hierarchy of master transcription factors.

The primary transcriptomic response to inorganic NPK fertilization is expansive and heavily biased toward gene repression, anchored predominantly by members of the AP2/ERF transcription factor superfamily (Nakano, Suzuki et al. 2006, Licausi, Ohme-Takagi et al. 2013). Multimodal integration across bulk expression, scRNA-seq, and scATAC-seq chromatin accessibility identified two top-tier hub regulators: *EREBP-4-like* and an ethylene-responsive *ERF*. Among these, *EREBP-4-like* functions as the primary regulatory node, governing the largest downstream network of 289 target genes. Functional enrichment of this regulon demonstrates a strong concentration in cytochrome P450 activity, monooxygenase functions, and iron/heme-binding processes, pointing to a coordinated metabolic realignment. Comparison of inorganic NPK with a mixed organic/inorganic regime (OMNPK) demonstrates that while the core regulatory architecture remains conserved across TF families, specific regulators exhibit regime-dependent dynamics. Most notably, *WRKY24* reverses its expression direction between regimes, showing strong induction under inorganic NPK but marked repression under the mixed amendment, positioning it as a potential sensor for organic versus inorganic nutrient sources.

Cellular localization predictions derived from non-negative least squares (NNLS) deconvolution and cell-type marker enrichment reveal that transcriptional reprogramming is highly tissue-specific. Transcriptional activity shifts heavily toward the central vascular cylinder, specifically the stele, phloem, xylem, and the mature pith, while the relative transcriptional contribution of the epidermis declines (Marand, Chen et al. 2021, Ortiz-Ramírez, Guillotin et al. 2021). This cellularly driven structural prioritization reflects the fundamental physiological role of vascular tissues in driving nutrient loading, solute transport, and systemic long-distance signaling following fertilizer uptake. Methodological validation using a Scaden-style deep neural network ensemble further reinforced these spatial insights. The deep-learning deconvolution model independently corroborated the vascular shift identified by NNLS while successfully recovering rare, low-abundance cell types, including the endodermis, phloem, and pericycle, that linear models frequently collapse to zero.

While core TF families remain active across both treatments, the predicted key master regulators exhibit striking regime-dependent dynamics. Most notably, WRKY24, DREB1A, and NAC61 undergo directional change, with WRKY24 switching from strong induction under inorganic NPK to marked repression under the mixed amendment, highlighting its potential function as a marker for organic versus inorganic nitrogen sources. Furthermore, while both fertilizer regimes share a primary stele-biased signature, the mature cortex exhibits divergent responses by expanding its transcriptomic presence under mixed organic/inorganic amendment while contracting under inorganic NPK alone. Predictive network modeling using CellOracle validated these relationships. Simulated knockouts of six primary hub regulators resulted in an 82.6% to 100% collapse in the expression of their predicted activation targets, alongside the de-repression of target genes specifically within their resident vascular and pith cell types. This tissue-confined disruption confirms that the network operates as a coherent, causal system. The validity of the multi-evidence tiering strategy was further confirmed by simulating a knockout of *NAC61*, a Tier 2 factor possessing accessible chromatin but lacking detectable single-cell expression. Knocking out *NAC61* produced limited (51%) downstream signal propagation, confirming that open chromatin alone, without active cell-type expression, is insufficient for regulatory driving capacity.

Broadening the scope to the rhizosphere holobiont, shotgun metagenomic profiling demonstrated that chemical fertilization induces a strong shift in bacterial community composition toward *Bacillus* dominance. Although specific microbial functional pathways did not clear multiple-testing correction due to single assembled sample constraints, these metagenomic findings establish crucial taxonomic context for host–microbiome co-adaptation under high-nutrient inputs. From an agronomic standpoint, this multi-omics framework condenses over 1,000 broad DEGs down to high-priority candidate nodes for crop engineering. High-confidence Tier 1 regulators like EREBP-4-like and the phloem-restricted WRKY24 node represent prime candidates for gene editing or molecular breeding aimed at optimizing vascular transport and nitrogen use efficiency (NUE).

## Limitations and Strategic Future Directions

While these findings provide a detailed framework for fertilizer response in maize, several limitations highlight key avenues for future research. First, the RNA-Seq analysis relied on a discovery-scale sample design (*n* = 2 per regime). This limitation was mitigated by cross-validating results against independent public datasets, including the deficiency dataset GSE125417 and single-cell nitrate response dataset GSE183171. Future field and laboratory trials will incorporate robust design and biological replication to improve the analytical power. Second, the regulatory network edges were inferred using public reference single-cell atlases. Future studies should perform direct scRNA-seq and spatial transcriptomics on roots harvested directly under specific fertilization treatments. Finally, to move beyond *in silico* knockout simulations, experimental validation must prioritize CRISPR-mediated knockout or inducible perturbation of *EREBP-4-like* and *WRKY24*, combined directly with treatment-matched ChIP-seq or CUT&Tag chromatin profiling.

## 5. Conclusions

Inorganic NPK fertilization of maize roots induces a large, directionally biased transcriptional response that is predicted to be spatially concentrated in vascular and pith tissues and organized around a small set of prioritized TF regulators. AP2/ERF-family factors, particularly *EREBP-4-like* and an ethylene-responsive *ERF,* were predicted as major regulators with chromatin-level support; *WRKY24* provides a phloem-specific node linking fertilizer status to systemic nitrogen signaling, and contextual *NAC/DREB1A* factors extend the regulatory space to cortex, endodermis, and the meristem. The rhizosphere microbiome shares the same perturbed functional categories as the host, a correspondence that, while not a statistically enriched co-response in this single-run metagenome, nominates specific soil–plant metabolic axes for replicated follow-up. By resolving fertilizer responses to specific cell types and prioritized regulators, this integrative framework provides candidate targets for molecular breeding and agronomic strategies aimed at improving nitrogen use efficiency in maize.

## Author contributions

B.M. and A.G. conducted bioinformatics analyses, B.M performed the single cell RNA-Seq data analysis, A.G. conducted the metagenome analysis. J.H. performed the metagenome taxonomy and bulk RNA-Seq integration. B.M. coordinated the research program and oversaw the experimental work, J.H., A.G., and B.M. wrote the manuscript. All authors discussed the results, critically reviewed the manuscript and provided feedback.

## Data and materials availability

All data needed to evaluate the conclusions in the paper are present in the paper and/or the Supplementary Materials. Additional data related to this paper may be requested from the authors.

