## Supplemental Text for "Multi-Omics Integration Predicts Cell-Specific Gene Regulatory Response and Rhizosphere Dynamics in Maize Root Fertilizer Treatment"

**Running title:** Predicted Cell-specific regulome of fertilizer response in maize root

**Authors:** Jade Horcoff^1^, Anuradha Goswami^1^, Bharat Mishra¹*

### **Affiliations:**

¹Department of Biological Sciences, University of Notre Dame, Notre Dame, IN

**Supplemental Information.**

**Supplementary Figures S1–S6**

**Figure S1:** Fertilizer-response transcriptome changes in maize root.

**Figure S2:** Single cell deconvolution predicts fertilizer associated cell types in maize roots.

**Figure S3:** Regulome activity in maize root atlas.

**Figure S4:** TF scRNA dot plot for percentage expression and mean expression of TFs across 22 root cell types.

**Figure S5:** Per-regulator regulome for the four remaining regulators.

**Figure S6:** Temporal organization of the fertilizer-responsive transcriptome along the diffusion pseudotime.

Supplementary Table S1-S4

**Table S1:** Maize Root RNA-Seq and GRN during fertilization treatment.

**Table S2:** Single cell deconvolution of fertilizer induced maize roots.

**Table S3:** Maize Root regulome activity.

**Table S4:** Maize rhizosphere metagenome in inorganic vs organic treatment.

**Supplementary Figures**
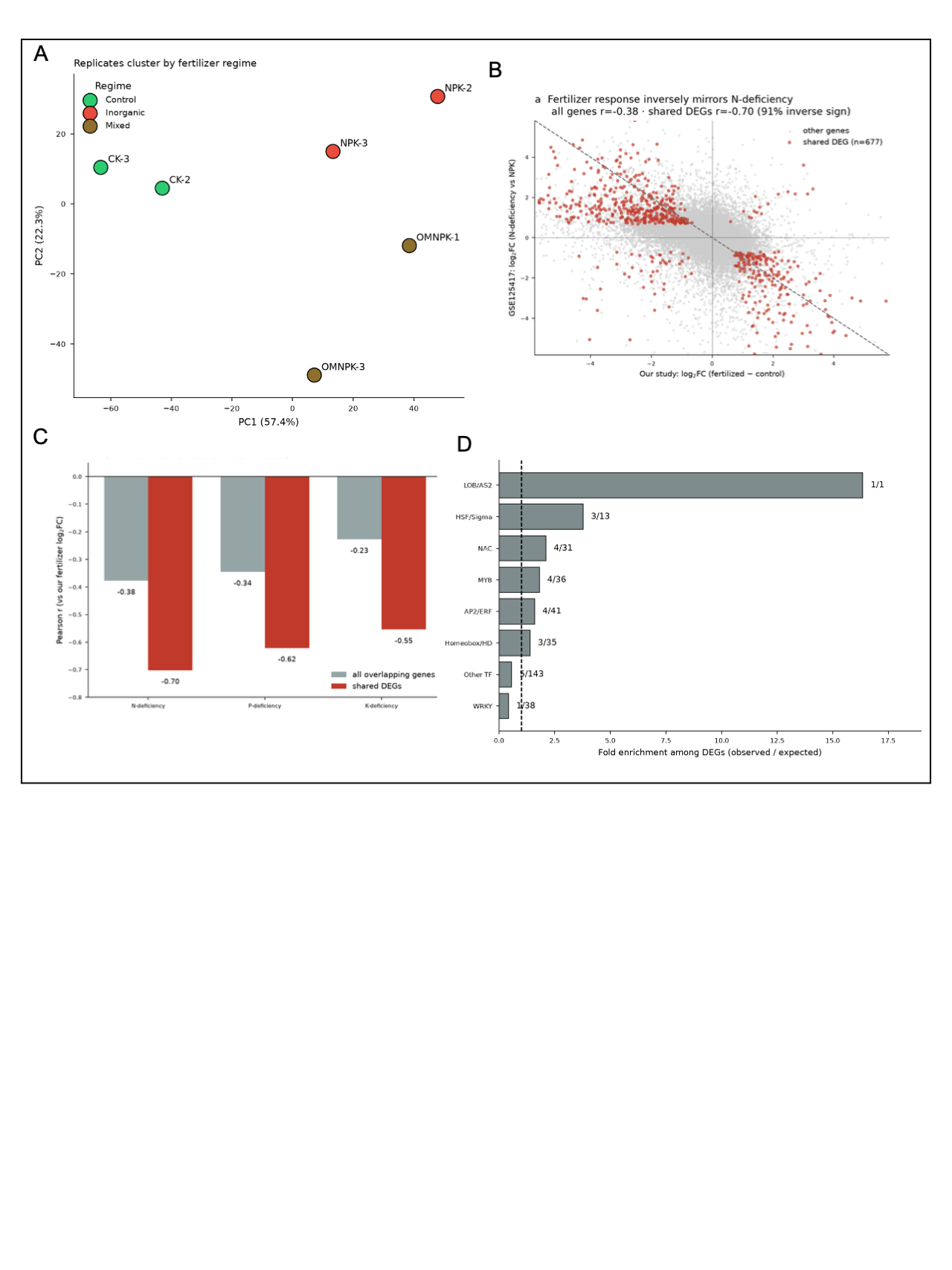

**Figure S1:** Fertilizer-response transcriptome changes in maize root. **(A)**Principal-component analysis of the six RNA-seq libraries (log₂ CPM of the 2,000 most variable genes), colored by fertilizer regime. PC1 (57.4% of variance) separates fertilized from control roots and replicates of each regime cluster together (between-group/within-group distance ratio 1.69), supporting the robustness of the differential-expression contrast despite two replicates per regime. **(B)** Gene-level comparison of our fertilizer response (log₂FC fertilized − control) against the GSE125417 nitrogen-deficiency response (log₂FC N-deficient vs NPK); grey = all overlapping genes, red = genes significant in both studies; the dashed anti-diagonal marks perfect inverse agreement. **(C)** Pearson correlation of our fertilizer log₂FC against each GSE125417 deficiency contrast, for all overlapping genes and for shared DEGs; the inverse concordance is strongest for nitrogen, identifying the NPK response as N-dominated. **(D)** Family-size-normalized TF enrichment among inorganic DEGs. Bars show fold-enrichment (observed/expected) per family; labels give DEGs/tested-members.

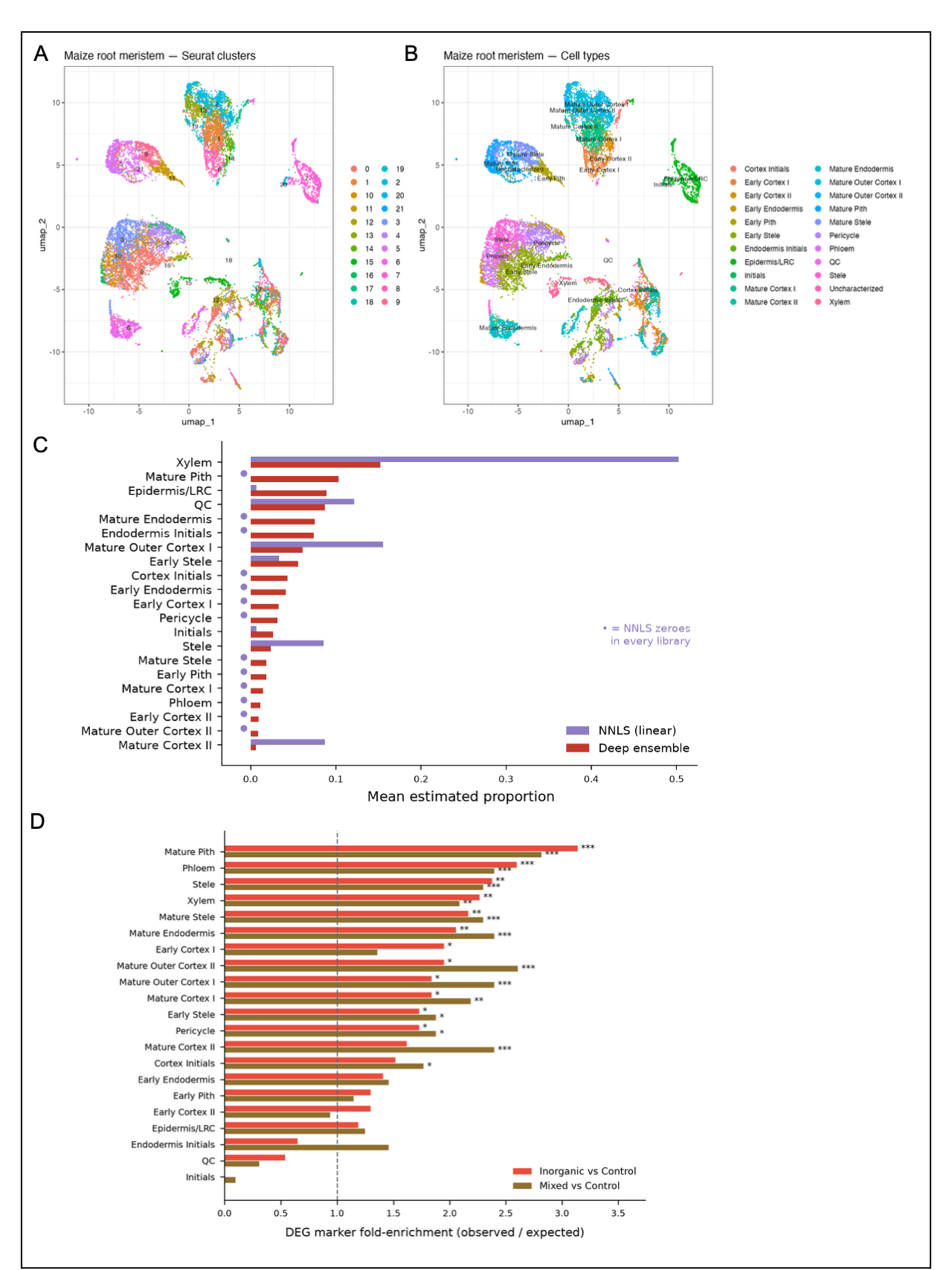

**Figure S2:** Single cell deconvolution predicts fertilizer associated cell types in maize roots. **(A-B)** UMAP of the maize root scRNA-seq atlas (GSE173087): Seurat clusters (left) and annotated cell types (right). **(C)** Mean estimated proportion per cell type across the six bulk libraries, NNLS (linear) versus the Scaden-style deep neural network ensemble. Cell types NNLS zeroes in every library are marked with a dot. **(D)** DEG cell-type marker fold-enrichment for the inorganic and mixed contrasts. Dashed line marks no enrichment (fold = 1); asterisks denote BH-adjusted significance (* < 0.05, ** < 0.01, *** < 0.001). Both contrasts converge on vascular/pith tissues; the mature-cortex classes are more strongly enriched under the mixed regime.

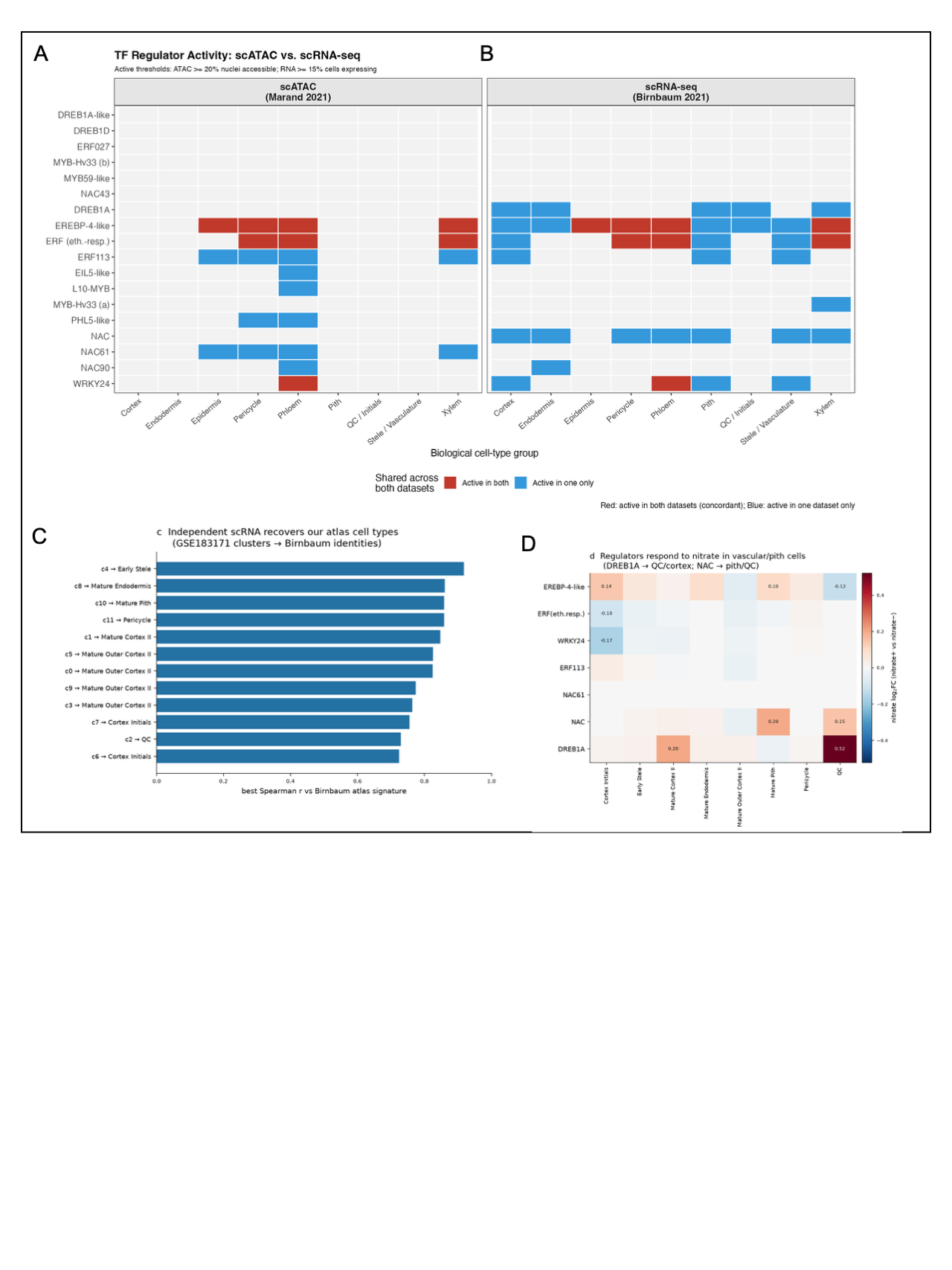

**Figure S3:** Regulome activity in maize root atlas. **(A-B)** TF chromatin accessibility from scATAC versus expression from scRNA per cell type. **(C)** GSE183171 single-cell clusters assigned to Birnbaum reference cell types by best Spearman correlation of mean expression against the atlas signature matrix. **(D)** Cell-type-resolved nitrate response (log₂FC nitrate+ vs nitrate−) of the seven prioritized regulators in GSE183171; DREB1A and NAC are induced in vascular/meristematic cell types (quiescent center, cortex, pith).

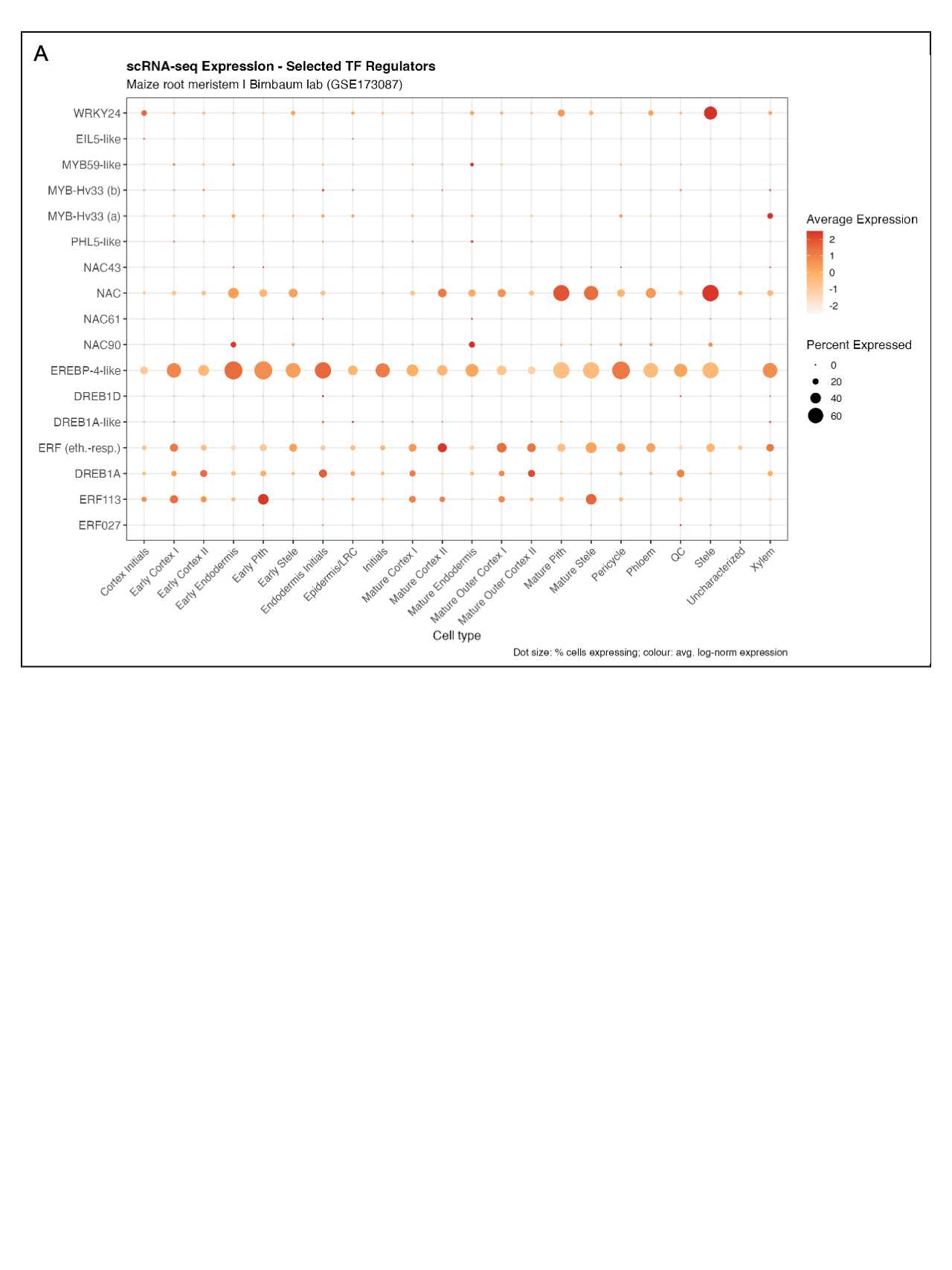

**Figure S4:** TF scRNA dot plot for percentage expression and mean expression of TFs across 22 root cell types.

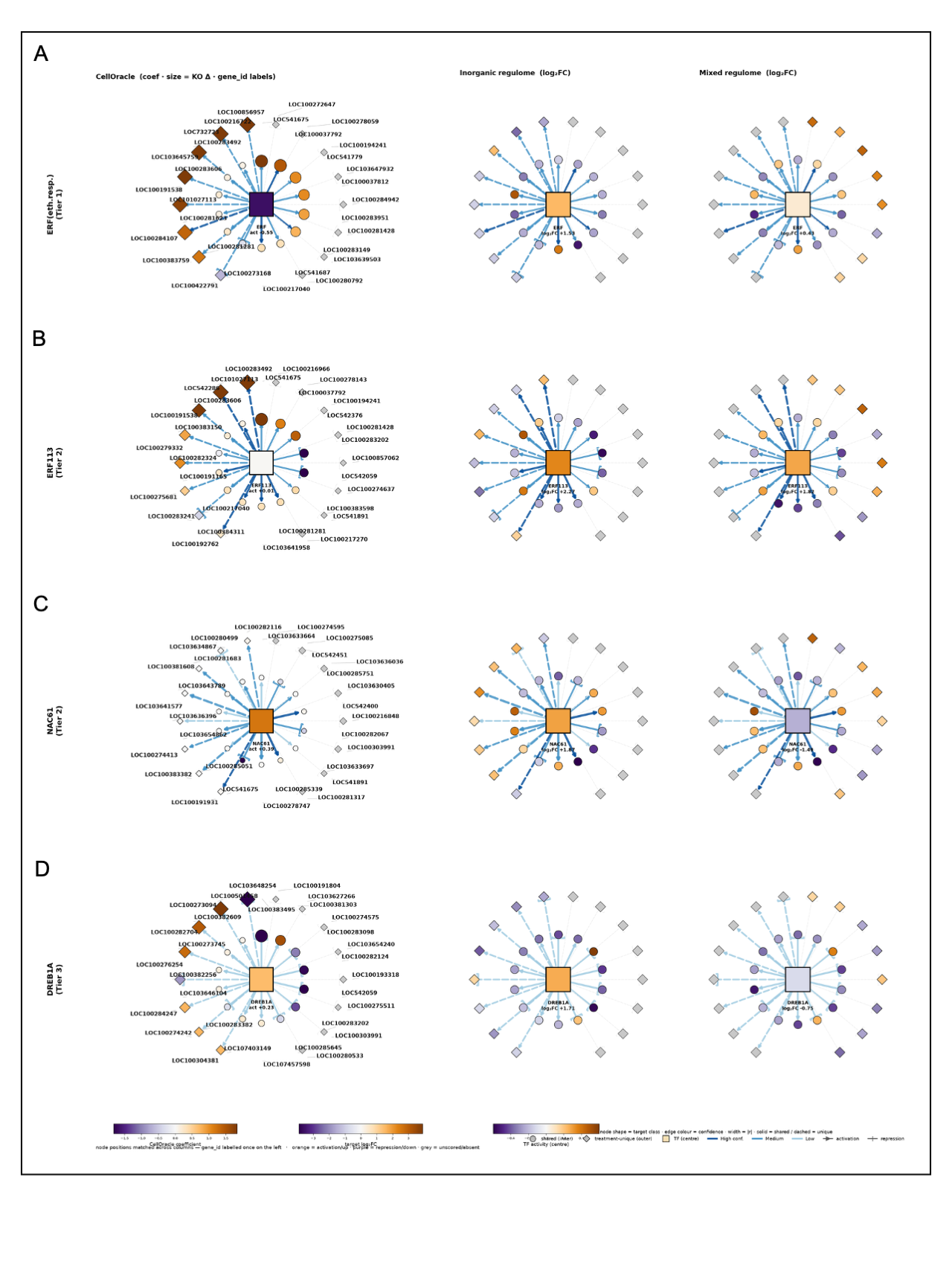

**Figure S5:** Per-regulator regulome for the four remaining regulators (ethylene-responsive ERF, ERF113, NAC61, DREB1A; A-D). Rows are regulators; columns show the CellOracle regulatory model (node color = coefficient, size = knockout Δ), the inorganic response and the mixed response (node color = target DESeq2 log₂FC). Circles = shared-core targets, diamonds = treatment-unique; solid/dashed edges = shared/unique links; arrowheads = activation, bars = repression; edge color = confidence, width = co-expression |r|; the central square is the regulator colored by its own activity. NAC61 and DREB1A show the same directional switch as WRKY24 (induced under inorganic NPK, repressed under the mixed amendment). Node positions matched across columns; gene identifiers labelled once on the CellOracle panel.

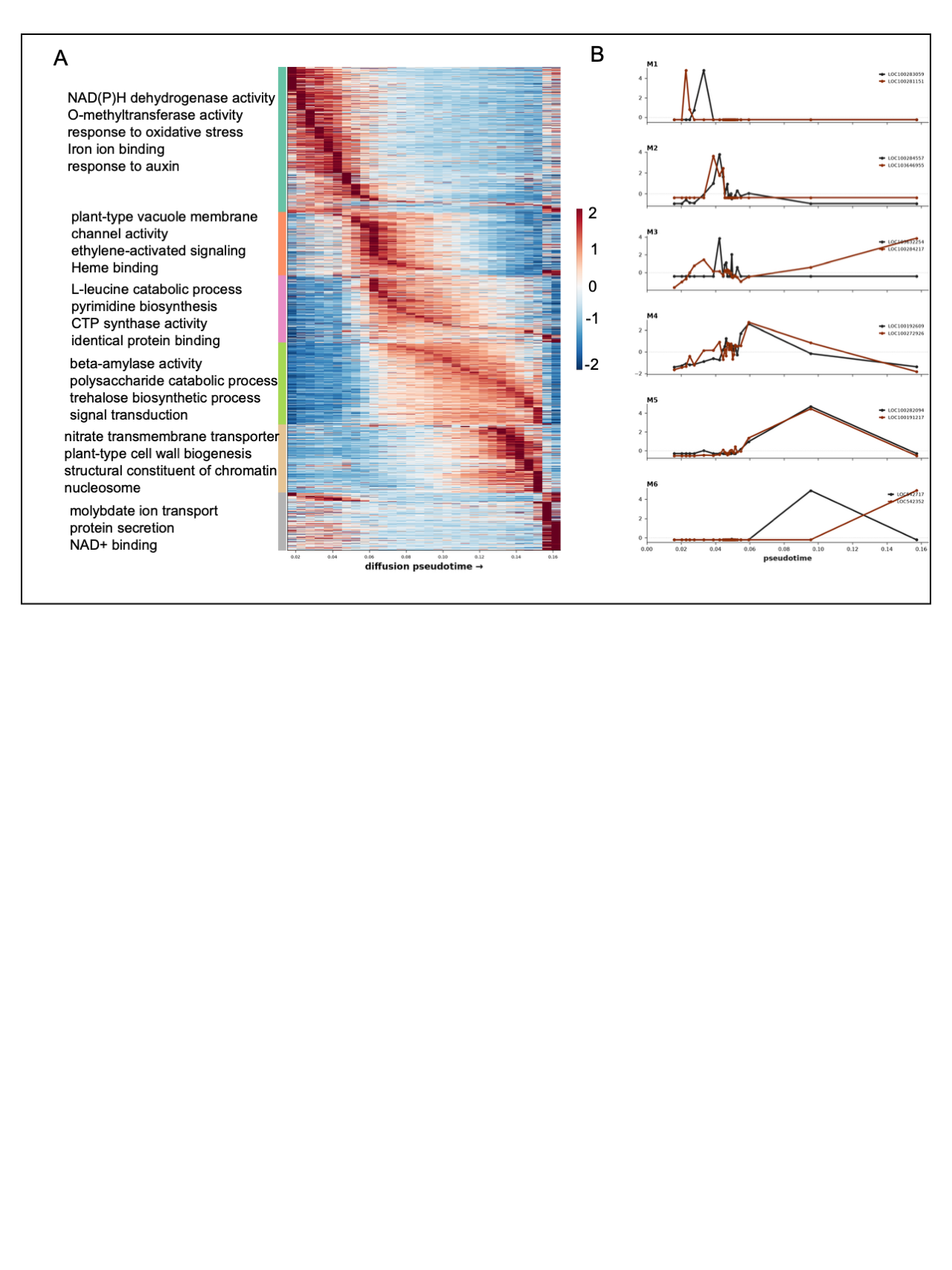

**Figure S6:** Temporal organization of the fertilizer-responsive transcriptome along the diffusion pseudotime. **(A)** Expression of the 1,848 union DEGs (rows) across 30 equal-cell pseudotime bins, z-scored per gene and hierarchically clustered (Ward) into six co-expression modules ordered by peak pseudotime. Left: Genome-wide GO enrichment per module (hypergeometric, BH-corrected *P* <0.05): early modules are dominated by cell-division and cell-wall machinery (microtubule, COPI vesicle coat, secondary cell wall biogenesis), progressing through transport and ethylene signalling to late modules enriched for nitrate transport, chromatin/nucleosome components and cellulose synthase. **(B)** Expression of the two most module-specific DEGs per module along pseudotime, binned into 25 equal-cell pseudotime windows and z-scored per gene to reveal the temporal pattern (raw single-cell values are dropout-dominated and uninformative).
